# PGG Potentiates Olaparib-Induced DNA Damage and STING-Associated Immunogenic Signaling to Enhance PD-L1 Blockade in Triple-Negative Breast Cancer

**DOI:** 10.64898/2026.09.26.752069

**Authors:** Sihan Liu, Xuan Li, Xiaochen Yang, Meiling Xiao, Tiancheng Fan, Jie Yu, Jie Zeng

**Author notes:** Correspondence: Jie Yu, 18670323423, Jie Zeng, 13974837749. These authors made equal contribution to this work and should be considered co-first authors (Sihan Liu, Xuan Li).

## Abstract

Triple-negative breast cancer (TNBC) is a highly aggressive malignancy and options for targeted therapy are limited. Although PARP inhibitors (PARPi) like olaparib (Ola) leverage synthetic lethality, their clinical utility is severely impeded by drug resistance and low frequency of BRCA mutations. In this study, 1,2,3,4,6-penta-O-galloyl-β-D-glucose (PGG), a natural polyphenolic compound, can induce a state of homologous recombination deficiency (HRD) in TNBC cells by inhibiting the PALB2-BRCA2 interaction. PGG potentiated the inhibitory effects of Ola on the in vitro proliferation, clonogenic potential, migration, and invasion of TNBC cells. Mechanistically, the dual therapy triggered severe oxidative stress and mitochondrial collapse, which accelerated the classical apoptosis cascade and promoted the emission of immunogenic cell death (ICD)-associated DAMPs. The accumulated cytosolic double-stranded DNA activated the innate immune cGAS-STING pathway, thereby promoting robust intra-tumoral infiltration of CD8^+^ T cells. Furthermore, since cGAS-STING hyperactivation drives compensatory PD-L1 upregulation on TNBC cells, incorporating anti-PD-L1 antibody into the combination regimen further reversed the immunosuppressive barrier, achieving near-complete tumor eradication and maximum infiltration of cytotoxic T cells in vivo. Bioinformatics analyses further suggested a potential network of PGG targets associated with extracellular matrix organization, offering a bioinformatics-derived hypothesis for its theoretical involvement in modulating stromal tension and vascular homeostasis. Collectively, this study establishes PGG as a multi-functional therapeutic agent integrating intracellular synthetic lethality and innate immune sensing, while providing a preliminary bioinformatic rationale for prospective microenvironmental investigation. This study also showes PGG as a multi-functional therapeutic agent that integrates intracellular synthetic lethality, innate immune sensing, and microenvironmental remodeling to convert immunologically “cold” TNBC into “hot” tumors.

## Introduction

Triple-negative breast cancer (TNBC) is a highly aggressive malignancy occupying approximately 15% to 20% of all breast cancer (BC) cases [1]. The insufficient estrogen receptor (ER), progesterone receptor (PR), and HER-2 render hormone therapy or HER-2-targeted therapy largely ineffective, leading to poor prognosis [2, 3]. Therefore, primary treatment strategies for TNBC are adjuvant and neoadjuvant chemotherapy [4]. Nevertheless, patients suffering advanced TNBC still exhibit poor outcomes alongside high mortality. Besides, a significant portion of patients show no response to standard chemotherapy, necessitating the development of novel therapeutic options.

Poly (ADP-ribose) polymerase 1 (PARP1) is a nuclear enzyme capable of recognizing and binding to DNA single-strand breaks, and subsequently recruiting factors to trigger the repair process by catalyzing the ADP-ribosylation of target proteins. PARP inhibitors (PARPi), including talazoparib (TA) and olaparib (Ola), demonstrate large promise in treating BC [5–9]. For instance, Ola specifically targets and kills BRCA1- or PTEN-deficient cells via synthetic lethality, thus providing a new paradigm for targeted therapy [10, 11]. Nevertheless, PARPi exhibits a limited clinical efficacy due to drug resistance and the low frequency of mutations in BC patients. Therefore, the current focus in BC research is to explore combination therapies to sensitize BC cells to PARPi.

Programmed cell death ligand 1 (PD-L1) is an immune checkpoint molecule expressed on tumor cells, capable of interacting with its ligand, PD-1 [12]. The PD-1/PD-L1 interaction constrains T cell activation and induces T cell exhaustion, restricting cytotoxic cytokine production and dampening the anti-tumor immune response [13]. Thus, T cells can exert an augmented anti-tumor response following the therapeutic blockade of the PD-1/PD-L1 axis [14]. Monoclonal antibodies designed to disrupt this pathway, collectively termed immune checkpoint inhibitors (ICIs), have dramatically reshaped the treatment paradigm for multiple solid tumors [15–17]. Among breast cancer subtypes, TNBC is generally considered a prime candidate for ICI regimens due to its relatively higher mutational burden and baseline immune signature [18]. Clinical outcomes, however, highlight a profound therapeutic heterogeneity; a significant proportion of patients derive negligible benefit from ICIs due to insufficient functional TILs, an active immunosuppressive milieu, or adaptive PD-L1 upregulation that collectively maintain a functionally “cold” TME. Overcoming this resistance requires combinatorial strategies capable of robustly stimulating systemic anti-tumor immunity [19, 20]. In this regard, exploiting synthetic lethality represents an attractive approach to trigger the release of endogenous danger signals and reactivate innate immune sensing within the tumor niche.

Natural products and their derivatives exhibit biocompatibility and pleiotropic mechanisms, making them valuable candidates for BC treatment. Flavonoid- and polyphenol-based drugs have demonstrated synergistic anti-tumor effects in BC [21]. For instance, PGG, a naturally occurring polyphenolic compound widely found in medicinal herbs of *Rhus chinensis* and *Paeonia suffruticosa*, exhibits potent anti-cancer efficacy [22, 23] with negligible cytotoxicity toward normal cells. Recently, PGG was identified as a specific small-molecule inhibitor of PALB2-BRCA2 interaction [24]. By directly binding to the PALB2 WD40 domain, PGG obstructs BRCA2 recruitment and the subsequent assembly of RAD51 recombinase at DNA double-strand breaks (DSBs), which selectively suppresses homologous recombination repair. This pharmacologically induced state of homologous recombination deficiency (HRD) produces robust synthetic lethality when combined with Ola, which not only reduces the *IC*_50_ values in vitro but also inhibits BRCA-wild-type BC patient-derived xenografts (PDXs) in vivo.

Cyclic-GMP-AMP synthase (cGAS) is a cytosolic innate immune sensor interacting with the sugar-phosphate backbone of double-stranded DNA (dsDNA) via positively charged amino acid residues [25]. Furthermore, activated cGAS works on synthesizing cGAMP in an ATP/GTP-dependent manner, subsequently binding to the immune adaptor protein STING and promotes its trafficking from the endoplasmic reticulum to the Golgi apparatus [26]. An obvious correlation was detected between the activation status of cGAS-STING pathway and HRD, high tumor mutational burden, and somatic copy number variations in TNBC [27]. STING enhances anti-tumor immune function through facilitating type I interferon and inflammatory cytokines to be largely produced, in turn accelerating the release of chemokines of CCL5 and CXCL10 [28]. This cascade subsequently drives cytotoxic CD8^+^ T cells, CD4^+^ T cells and CD20^+^ B cells to be infiltrated into the TME, thus effectively bridging innate and adaptive immunity [29]. Therefore, the cGAS-STING pathway presents a valuable target for alleviating immune suppression in breast tumors and remodeling the TME into an immunologically “hot” state, thereby enhancing tumor responsiveness to the combination of ICIs with chemotherapy or PARPi.

Based on these insights, we rationalized that the capacity of PGG to disrupt the PALB2–BRCA2 interaction and phenocopy an HRD state—an established mechanistic premise from prior literature [24], which could be leveraged to fuel downstream immune pathways. While this pharmacologically induced HRD serves as the foundational hypothesis of our study, it remains to be elucidated whether the resulting DNA lesions can actively drive the cGAS–STING network to reshape the TME and reactivate anti-tumor immunity in TNBC. According to our finding, PGG activates the STING signaling pathway in TNBC cells and augments the therapeutic efficacy of PARPi, while simultaneously orchestrating immune modulation via the upregulation of PD-L1. Furthermore, PGG-induced PD-L1 upregulation also enhanced the therapeutic efficacy of anti-PD-L1 antibody (aPD-L1) in syngeneic mouse models. Collectively, all these elucidate the molecular mechanisms pertaining to PGG-mediated sensitization of TNBC cells to PARPi and PD-L1 induction, thus paving the way for novel combination strategies tailored for patients suffering PD-L1^+^ TNBC.

## Materials and Methods

### Data Sources

Potential targets of PGG were retrieved and downloaded from the HERB database (http://herb.ac.cn), an inventory of traditional Chinese medicine (TCM) formulations and their chemical constituents, using ‘1,2,3,4,6-O-pentagalloylglucose’ as the search query. Concurrently, the disease-related target genes for TNBC were screened from the GeneCards database (https://www.genecards.org) using the keywords ‘triple negative breast cancer’. Finally, the overlapping target genes between PGG and TNBC were identified and visualized using a Venn diagram generated via Visual Paradigm.

### GO Functional Annotation and KEGG Pathway Enrichment Analysis

The common targets of PGG and TNBC underwent GO and KEGG enrichment analyses via Metascape (https://metascape.org). In the GO analysis, the top 10 enriched terms each from the biological process (BP), cellular component (CC), and molecular function (MF) categories were retrieved to construct a bubble plot. The pharmacological pathways of PGG were identified through KEGG analysis; the enriched pathways were screened with a threshold of *P* < 0.05, and visualized using a dot plot.

### Cell Culture

Human (MDA-MB-231) and murine (4T1) TNBC cell lines underwent culture in Dulbecco’s Modified Eagle Medium (DMEM) supplemented with 10% fetal bovine serum (FBS) and 1% penicillin-streptomycin (PS) solution in a humidified incubator at 37°C with a 5% CO_2_ atmosphere. All cell lines were authenticated via STR profiling, confirmed negative for mycoplasma contamination, and restricted to passages below 15 for all experiments.

### CCK-8 Assay

The 4T1 and MDA-MB-231 cells were seeded into 96-well plates at a density of 3×10³ cells/well and 5×10³ cells/well, respectively, with six replicate wells configured for each experimental group. The peripheral wells were filled with 100 μL of sterile PBS to eliminate the edge effect. Following cell adherence, the medium was replaced with fresh complete medium containing PGG (1 μM), Ola (10 μM), or their combination for 24, 48, or 72 h. At each specified time point, 10 μL of CCK-8 reagent was added into each well, followed by incubation at 37℃ for 1 h in the dark. The absorbance at 450 nm was measured using a microplate reader to evaluate cell viability. All experiments were performed in triplicate independently.

### Colony Formation Assay

TNBC cells were seeded into 6-well plates at a density of 500 cells/well and allowed to adhere overnight, and then cultured in the presence of DMSO, PGG (1 μM), Ola (10 μM), or PGG+Ola for 24 h. The treatment medium was subsequently replaced with fresh complete medium, which was replenished every 3 days. After 7–10 days of incubation, the visible colonies were fixed with 4% paraformaldehyde (PFA) for 15 min and stained with 0.1% crystal violet for another 15 min. Surviving colonies containing more than 50 cells were photographed and counted. Each group was assessed in triplicate, and the assay was repeated three times independently.

### Transwell Migration and Invasion Assays

Cell motility and invasiveness were evaluated with 24-well Transwell chambers (8-*μ*m pore size). For the migration assay, the suitably treated 4T1 and MDA-MB-231 cells were harvested, resuspended in 200 *μ*L serum-free DMEM, and seeded into the upper chambers. For the invasion assay, the upper chambers were pre-coated with Matrigel before cell seeding. The lower chambers were filled with 600 *μ*L complete medium containing 10% FBS as a chemoattractant. Subsequent to 24 h of incubation, experimenters carefully removed the cells that failed to traverse the membrane using a cotton swab. The migrated or invaded cells attached to the lower surface of the membrane underwent fixation with 4% PFA and 0.1% crystal violet staining in sequence. An optical microscope was employed for capturing representative multi-field images, alongside the counting of the cells.

### Measurement of Intracellular Reactive Oxygen Species (ROS)

Intracellular ROS production was monitored with a DCFH-DA staining kit. In brief, 4T1 and MDA-MB-231 cells were cultured in 6-well plates, followed by 24 h of treatment with the different agents. After PBS wash, the cells were incubated with 10 μM DCFH-DA probe diluted in serum-free medium at 37°C for 20 min in the dark. Following subsequent washes to remove extracellular dye, the cells were harvested, resuspended in PBS, and immediately analyzed by flow cytometry using the FITC channel. At least 10,000 events were collected per sample to quantify the mean fluorescence intensity (MFI) of DCF.

### Mitochondrial Membrane Potential (MMP) Analysis

The JC-1 probe was employed for MMP evaluation according to the manufacturer’s instructions. The suitably treated TNBC cells were incubated with the JC-1 working solution at 37°C for 20 min in the dark. Following two washes using ice-cold JC-1 staining buffer, the cells were observed and imaged under a fluorescence microscope. To ensure objective quantification, at least five randomized microscopic fields were captured per group using identical exposure settings. The fluorescence intensities of both the green (monomer) and red (aggregate) channels were subsequently measured using ImageJ software, and the relative green-to-red fluorescence ratio was calculated to evaluate mitochondrial depolarization.

### Cell Transfection and Western Blotting

For transient knockdown of STING, TNBC cells were transfected with STING-targeted siRNA (siSTING) or a non-targeting negative control siRNA (siNC) using Lipofectamine 3000 (Invitrogen) according to the manufacturer’s protocol. At 24 h post-transfection, cells were harvested, re-seeded into 6-well plates, and exposed to DMSO, PGG (1 μM), Ola (10 μM), or their combination for an additional 24 h. This yielded eight distinct experimental cohorts: (1) siNC + DMSO, (2) siNC + PGG, (3) siNC + Ola, (4) siNC + PGG + Ola, (5) siSTING + DMSO, (6) siSTING + PGG, (7) siSTING + Ola, and (8) siSTING + PGG + Ola. Subsequent to washing with ice-cold PBS, the cells underwent 30 min of lysis treatment on ice using RIPA buffer. Cells underwent centrifugation for the collection of supernatants, and a BCA assay kit was applied to the quantification of protein level. Equal amounts of protein received denaturation before SDS-PAGE separation, and were electro-transferred onto activated PVDF membranes using a wet transfer system. Following overnight incubation with primary antibodies against STING, p-STING, IRF3, p-IRF3, PD-L1, and vinculin, the membranes received certain period of incubation with HRP-conjugated secondary antibodies. An ECL detection system was employed for visualizing the protein bands, and the gray value ratios relative to the internal control (vinculin) were quantified by employing ImageJ software to assess relative protein expression levels.

### Syngeneic Tumor Modeling and Treatment Regimen

Female BALB/c mice (7-8 weeks old, weighing 20 ± 2 g) were inoculated subcutaneously in the right flank with 1 × 10^6^ 4T1 cells to establish subcutaneous tumors. With the tumor volumes reaching ∼50 mm^3^ (day 8), the mice fell into four treatment cohorts: DMSO control, Ola (50 mg/kg, i.p., q.d.), PGG (10 mg/kg, i.v., q.2.d.), and PGG+Ola. In another experiment, the treatment groups were IgG control, aPD-L1 (10mg/Kg, i.p., q.d.), Ola+aPD-L1, PGG+aPD-L1, PGG+Ola and PGG+Ola+aPD-L1. Tumor dimensions were recorded every two days, and the volumes were estimated by the formula V = (L×W^2^)/2. After the euthanasia of mice at the experimental endpoint (day 16), their tumors were excised, weighed, and fixed in 4% PFA. All animal experimental procedures strictly followed the guidelines of the Experimental Animal Ethics Committee of Hunan Normal University (Approval No. [2024] 97).

### Histological Analysis and Immunohistochemistry

Subsequent to 24 h of fixation in 4% PFA, harvested tumor tissues underwent ethanol dehydration, xylene clearing and paraffin-embedding in sequence, followed by being sectioned into 5-μm slices using a microtome. For histopathological evaluation, representative sections underwent standard H&E staining for the assessment of morphological changes and tissue necrosis. For immunohistochemical analysis, the sections were deparaffinized before rehydration, and then received heat-induced antigen retrieval in a 10 mM citrate buffer (pH 6.0) using a microwave. 3% hydrogen peroxide was employed for quenching the endogenous peroxidase activity for 10 min, followed by 1 h of blockage of non-specific binding sites using 5% bovine serum albumin (BSA) at room temperature (RT). The sections then received one night of incubation at 4°C with a primary antibody against Ki-67. Subsequent to PBS wash, the tissues underwent 1 h of incubation with a HRP-conjugated secondary antibody at RT. Immunoreactivity was visualized by employing a 3,3’-diaminobenzidine (DAB) substrate kit, followed by hematoxylin counterstaining. An optical microscope was used for capturing relevant images, and the percentage of Ki-67-positive proliferating cells was quantified by counting positive nuclei relative to total nuclei in five randomly selected fields per section via ImageJ software.

### TUNEL Assay

A TUNEL apoptosis detection kit was adopted for evaluating the apoptosis in tumor tissues as per the producer’s protocol. In brief, 5-*μ*m-thick paraffin sections were rehydrated after deparaffinization, and then underwent 20 min of permeabilization with proteinase K (20 *μ*g/mL) at RT to facilitate enzyme penetration. After PBS wash, the sections underwent 1 h of incubation with the TUNEL reaction mixture encompassing terminal deoxynucleotidyl transferase (TdT) and fluorescein-dUTP in a humidified chamber at 37 ℃ in the dark. Nuclei then underwent 5 min of DAPI counterstaining to delineate the cellular architecture. The sections were mounted using an anti-fade mounting medium and confocal laser scanning microscopy served for relevant visualization. Apoptotic cells were identified by green fluorescence, and the apoptotic index was determined as the ratio of TUNEL-positive cells to the total number of DAPI-stained nuclei across multiple randomized microscopic fields.

### Immunofluorescence

For immunofluorescence staining of the TME, paraffin sections were rehydrated after deparaffinization, followed by the heat-induced antigen retrieval in EDTA buffer (pH 9.0). To minimize non-specific background fluorescence, sections underwent 1 h of incubation with 10% normal goat serum containing 0.3% Triton X-100 at RT. The sections then received one night of incubation at 4°C with primary antibodies targeting CD8 or PD-L1. After rigorous washing using PBST (PBS containing 0.1% Tween-20), the slides were subjected to 1 h of incubation with corresponding fluorophore-conjugated secondary antibodies at RT in the dark. Nuclei underwent 5 min of DAPI counterstaining. Fluorescence profiles were examined and captured under a confocal laser scanning microscope using identical exposure and gain settings across all groups to ensure comparability. Quantitative analysis was conducted using ImageJ software by calculating the MFI of CD8 and PD-L1 signals from ≥ 5 independent, randomly selected fields per specimen.

### Statistical Analysis

Statistical analyses were performed using GraphPad Prism 10.0 and SPSS 27.0. Quantitative datasets obtained from ImageJ analysis or experimental measurements are expressed as the mean ± standard error of the mean (SEM). The normality of data distribution was evaluated using the Shapiro-Wilk test, and homogeneity of variance was assessed via Levene’s test. For multi-group comparisons at a single clinical or experimental endpoint, one-way analysis of variance (ANOVA) followed by Tukey’s post-hoc test was performed. Longitudinal dynamic datasets, specifically in vitro cell proliferation and in vivo dynamic tumor growth curves, were analyzed using two-way repeated-measures ANOVA with Tukey’s multiple comparisons test. Bioinformatic enrichment analyses (GO and KEGG) were evaluated based on hypergeometric tests with Benjamini-Hochberg correction for false discovery rate (FDR < 0.05 or adjusted *P* < 0.05). Statistical significance was defined as *P* < 0.05 (* *P* < 0.05, ** *P* < 0.01, *** *P* < 0.001).

## Results

We identified 63 overlapping target genes between the 115 PGG-associated targets retrieved from the HERB database and 10,747 TNBC-related targets from GeneCards, aiming to elucidate the potential molecular mechanisms of PGG against TNBC (**Figure 1A**). To organically understand their functional networks, these common targets underwent GO and KEGG enrichment analyses. According to GO annotation, the intersecting targets were predominantly clustered in biological processes and molecular functions related to extracellular matrix (ECM) disassembly, humoral immune response, and endopeptidase activities (**Figure 1B**). Consistently, KEGG pathway analysis highlighted the obvious enrichment in the complement and coagulation cascades, transcriptional misregulation in cancer, and the renin-angiotensin system (**Figure 1C**). Taken together, these bioinformatic insights suggest that PGG may exert its anti-TNBC efficacy through a multi-target network, characterized by the dual capacity to disrupt oncogenic transcription within cancer cells and potentially modulate the immuno-coagulation crosstalk and vascular microenvironment within the tumor stroma.

**Figure 1.**
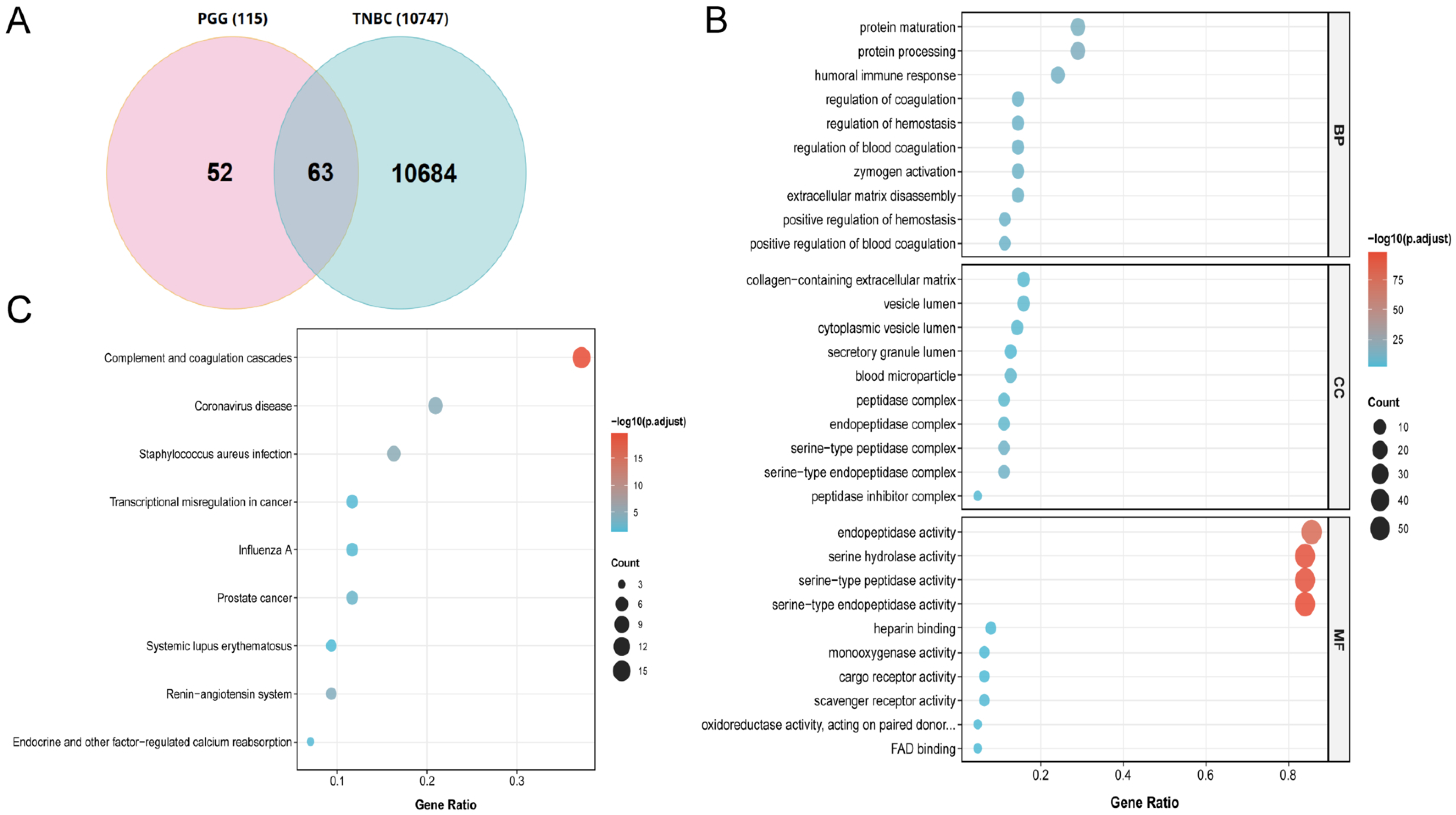
Identification and functional annotation of overlapping targets between PGG and TNBC. **(A)** Venn diagram showing 63 common target genes of PGG (115 targets from the HERB database) and TNBC (10,747 targets from the GeneCards database). **(B)** Bubble plot demonstrating the significantly enriched GO terms for the intersecting target genes. **(C)** Bubble plot showing the significantly enriched KEGG pathways involved in the therapeutic mechanisms of PGG against TNBC.

For more deeply elucidating the precise molecular mechanisms pertaining to the synergy of PGG with DNA-damaging agents, we conducted a target-specific enrichment analysis focusing on DNA damage response, ROS metabolism, and anti-tumor immunity (**Figure 2**). Within this functional network, functional mapping revealed significant enrichment in stress-responsive and oxidative regulation pathways. Concurrently, a prominent cluster of immune-regulatory signatures emerged, dominated by lymphocyte-mediated adaptive immunity and cell surface receptor signaling. Furthermore, pathways governing extracellular matrix disassembly and cytolysis were closely integrated into this network, indicating an intrinsic crosstalk between tumor cell death and To investigate the molecular mechanisms underlying the potential synergy between PGG and Ola, we first evaluated the expression and activation profiles of the STING signaling pathway. Compared with the DMSO control, PGG monotherapy exerted negligible effects on the phosphorylation of STING and IRF3, as well as PD-L1 expression, in both cell lines. In contrast, Ola monotherapy significantly augmented p-STING and p-IRF3 levels, while concurrently upregulating PD-L1 expression. Remarkably, co-treatment with PGG and Ola further amplified p-STING, p-IRF3, and PD-L1 levels beyond the baseline induction achieved by Ola monotherapy (**Figure 4A–H**). These findings demonstrate that Ola acts as the important driver of STING activation, a phenomenon that is robustly potentiated by PGG. To explore whether the PGG and Ola combination triggers an ICD-like phenotype, we examined the cellular redistribution of key damage-associated molecular patterns (DAMPs), specifically CRT and HMGB1. While PGG or Ola monotherapy elicited minor CRT exposure, their combination significantly accelerated its translocation to the plasma membrane in both TNBC cell lines (**Figure 4I–K**). Concurrently, immunofluorescence analysis revealed that HMGB1, which was predominantly sequestered within the nuclei of control and PGG-treated cells, underwent robust nuclear efflux following PGG+Ola dual therapy (**Figure 4L–N**). Taken together, these data suggest that PGG works in concert with Ola to induce ICD-associated DAMP release in TNBC cells, potentially contributing to subsequent microenvironmental remodeling and adaptive immune activation.

**Figure 2.**
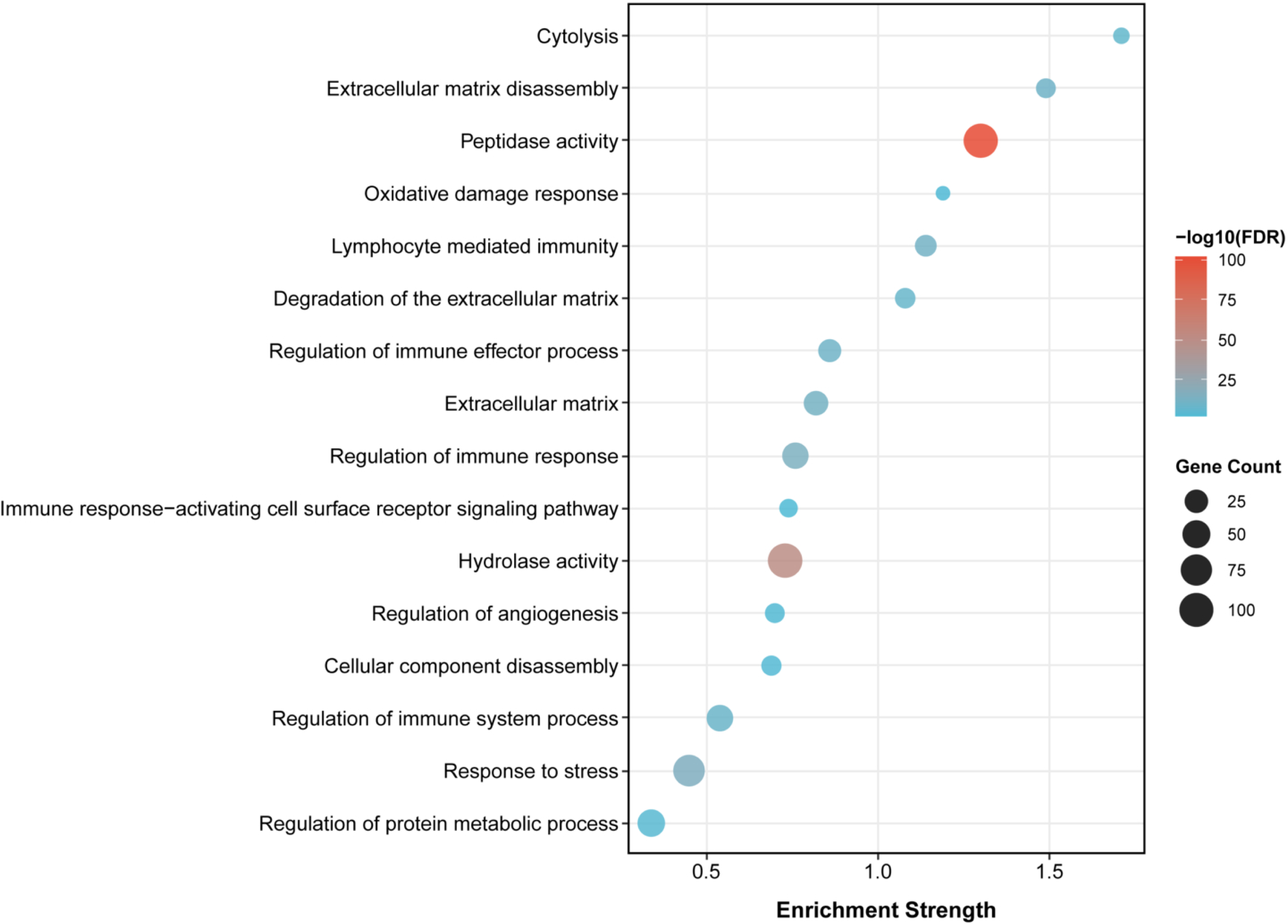
Target-specific enrichment analysis of PGG and TNBC intersection genes focusing on DNA damage, ROS metabolism, and immune regulation.

**Figure 3.**
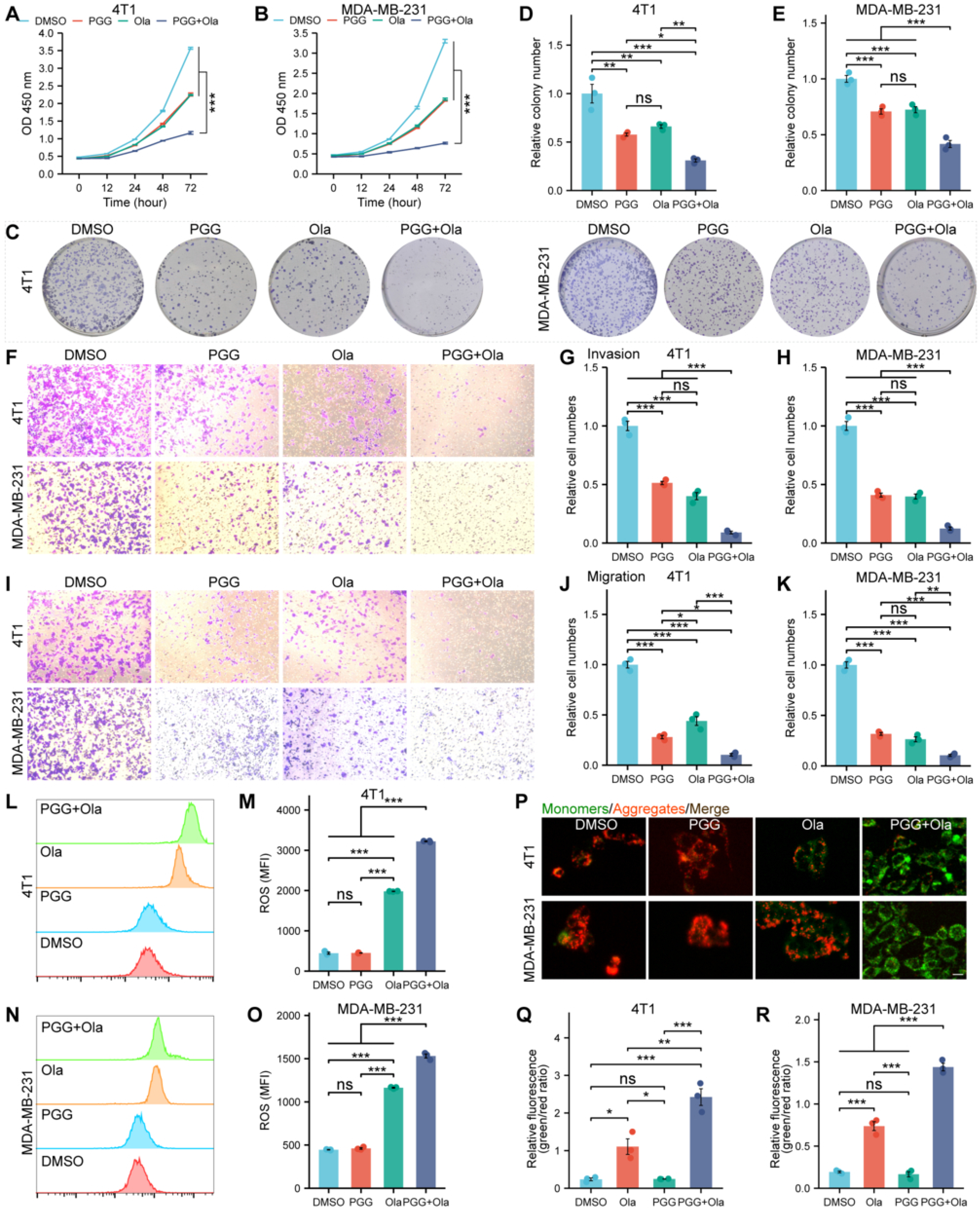
PGG and Ola exerted enhanced inhibition on TNBC cells in vitro by inducing oxidative stress and mitochondrial damage. **(A, B)** Viability rates of **(A)** 4T1 and **(B)** MDA-MB-231 cells at designated time points. **(C)** Representative images of colonies formed by the TNBC cells. **(D, E)** Number of colonies formed by the 4T1 **(D)** and MDA-MB-231 **(E)** cells after indicated treatments. **(F)** Representative transwell images showing the invasion of TNBC cells. **(G,H)** Quantification of the invasion capacities of 4T1 **(G)** and MDA-MB-231 **(H)** cells. **(I)** Representative transwell images showing the migration of TNBC cells. **(J, K)** Quantification of the migration capacities of 4T1 **(J)** and MDA-MB-231 **(K)** cells. **(L, N)** Flow cytometric histograms of 4T1 **(L)** and MDA-MB-231 **(N)** cells stained with DCFH-DA. **(M, O)** MFIs corresponding to intracellular ROS levels in 4T1 **(M)** and MDA-MB-231 **(O)** cells. **(P)** Representative fluorescence images illustrating JC-1 monomers (green), JC-1 aggregates (red), and merged signals in the indicated groups. **(Q, R)** Green/red fluorescence ratios corresponding to changes in MMP in the 4T1 **(Q)** and MDA-MB-231 **(R)** cells. Data are in the format of mean ± SEM. ns, not significant; \**P* < 0.05, \*\**P* < 0.01, \*\*\**P* < 0.001.

**Figure 4.**
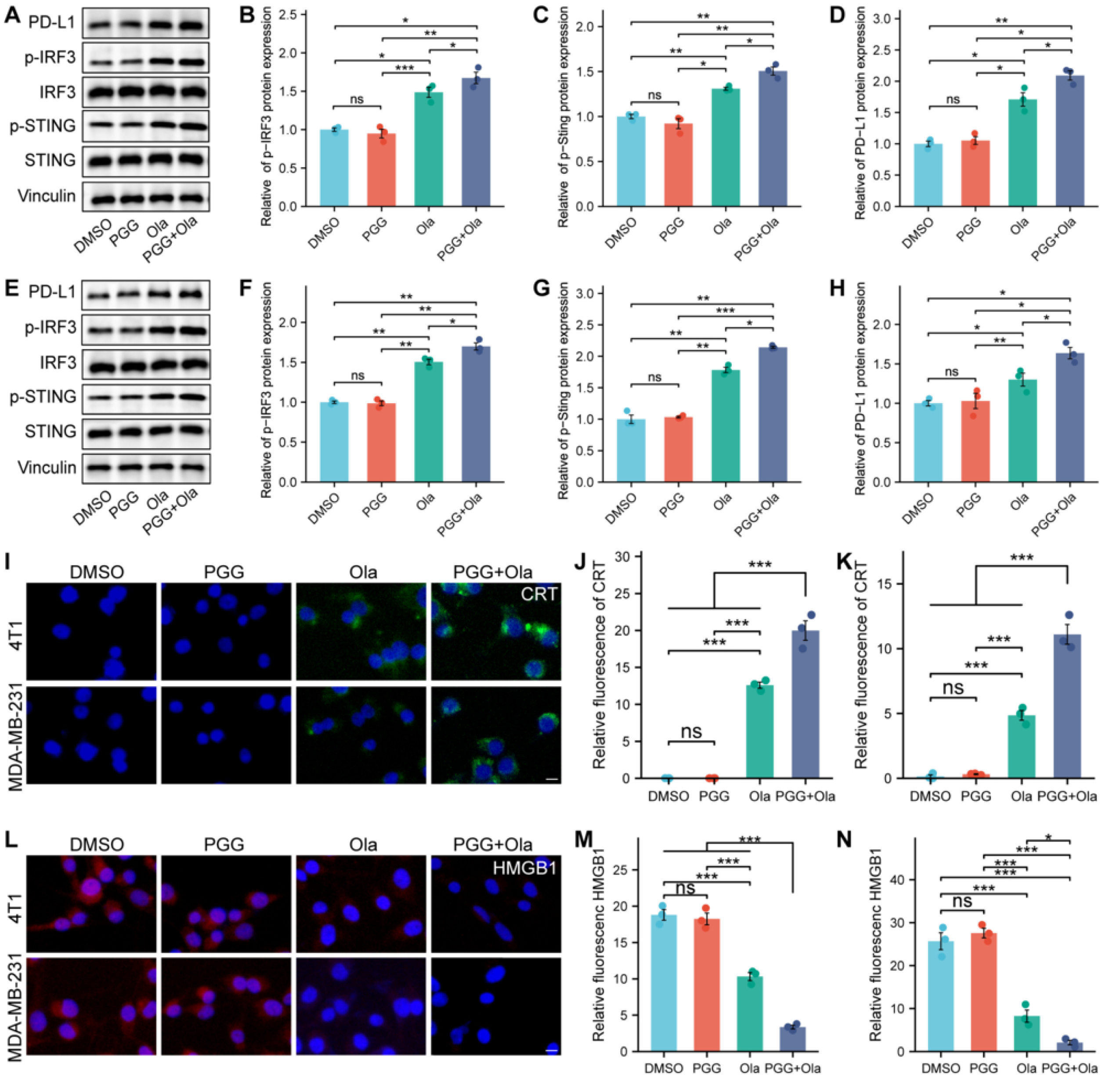
PGG and Ola activated the STING pathway, upregulated PD-L1, and triggered ICD-associated DAMP release in TNBC cells. **(A)** Representative immunoblots showing the bands for IRF3, p-IRF3, STING, p-STING, and PD-L1 in the 4T1 cells under the treatment of DMSO+siNC, PGG+siNC, Ola+siNC, and PGG+Ola+siNC. **(B-D)** Densitometric quantification of p-IRF3 **(B)**, p-STING **(C)** and PD-L1 **(D)** expression levels in the 4T1 cells. **(E)** Representative immunoblots showing the bands for IRF3, p-IRF3, STING, p-STING, and PD-L1 in the MDA-MB-231 cells under the treatment of DMSO+siNC, PGG+siNC, Ola+siNC, and PGG+Ola+siNC. **(F-H)** Densitometric quantification of p-IRF3 **(F)**, p-STING **(G)** and PD-L1 **(H)** expression levels in the MDA-MB-231 cells. **(I)** Representative immunofluorescence images illustrating the surface translocation of calreticulin (CRT, green) in 4T1 and MDA-MB-231 cells. Nuclei received DAPI counterstaining (blue). Scale bar = 20 μm. **(J, K)** Quantification of the relative fluorescence intensity of CRT in 4T1 **(J)** and MDA-MB-231 **(K)** cells. **(L)** Representative immunofluorescence images showing the intracellular localization and release of high mobility group box 1 (HMGB1, red) in 4T1 and MDA-MB-231 cells. Nuclei received DAPI counterstaining (blue). Scale bar = 20 μm. **(M, N)** Quantification of the relative fluorescence intensity of HMGB1 in 4T1 **(M)** and MDA-MB-231 **(N)** cells. Data are in the format of mean ± SEM. ns, not significant; \**P* < 0.05, \*\**P* < 0.01, \*\*\**P* < 0.001.

To determine whether the potential synergistic anti-tumor and DAMP-releasing effects of PGG and Ola are fundamentally mediated by STING activation, we performed genetic loss-of-function experiments via siRNA-mediated knockdown of STING (siSTING). As anticipated, STING silencing effectively abolished the PGG+Ola-induced phosphorylation of both STING and its downstream effector IRF3 in 4T1 and MDA-MB-231 cells (**Figure 5A–C, E–G**). Notably, while baseline PD-L1 expression remained partially elevated—likely driven by persistent upstream DNA damage—its upregulation by the combination therapy was significantly blunted upon STING depletion (**Figure 5D, H**). Consistently, STING depletion markedly diminished PGG+Ola-stimulated CRT plasma membrane translocation and completely blocked the nuclear efflux of HMGB1, causing its noticeable nuclear retention in both TNBC cell lines (**Figure 5I–N**). Collectively, these findings demonstrate that PGG potentiates Ola-mediated innate immune activation and DAMP emission in TNBC cells primarily through the activation of the STING signaling pathway.

**Figure 5.**
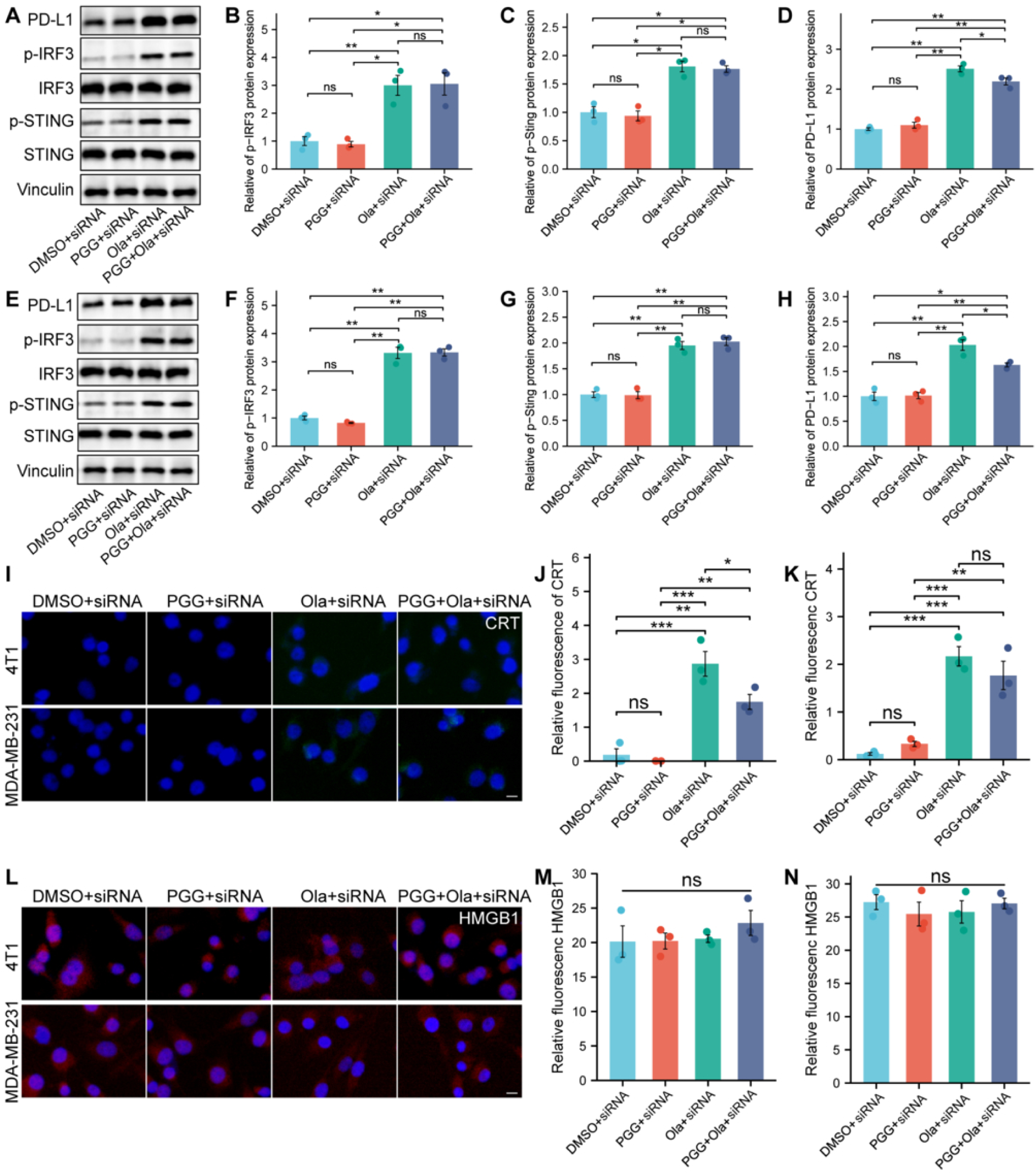
STING signaling mediates PGG- and Ola-induced innate immune activation and ICD-associated DAMP release in TNBC cells. **(A)** Representative immunoblots of IRF3, p-IRF3, STING, p-STING, and PD-L1 in 4T1 cells transfected with siSTING followed by treatment with DMSO, PGG (1 μM), Ola (10 μM), or their combination. **(B-D)** Densitometric quantification of p-IRF3 (B), p-STING (C) and PD-L1 (D) expression levels in the 4T1 cells. **(E)** Representative immunoblots of IRF3, p-IRF3, STING, p-STING, and PD-L1 in MDA-MB-231 cells under corresponding transfection and treatment regimens. **(F-H)** Densitometric quantification of p-IRF3, p-STING, and PD-L1 expression levels in the MDA-MB-231 cells. **(I)** Representative immunofluorescence images illustrating the surface translocation of CRT (green) in 4T1 and MDA-MB-231 cells following STING knockdown. Nuclei were counterstained with DAPI (blue). Scale bar = 20 μm. **(J, K)** Quantification of the relative surface fluorescence intensity of CRT in 4T1 (J) and MDA-MB-231 (K) cells. **(L)** Representative immunofluorescence images illustrating the intracellular localization and nuclear retention of HMGB1 (red) following STING knockdown. Nuclei were counterstained with DAPI (blue). Scale bar = 20 μ m. **(M, N)** Quantification of the relative intracellular fluorescence intensity of HMGB1 in 4T1 (M) and MDA-MB-231 (N) cells. Note: siRNA indicates STING-targeting siRNA, siSTING. Data are in the format of mean ±SEM. ns, not significant; \**P* < 0.05, \*\**P* < 0.01, \*\*\**P* < 0.001.

For assessing the anti-tumor activity of the PGG and Ola combination in vivo, an immunocompetent murine subcutaneous tumor model was established using 4T1 cells (**Figure 6A**). In **Figure 6D**, tumors in the DMSO control group progressed rapidly, reaching a volume close to 500 mm³ by day 16 post-inoculation. In contrast, mice treated with Ola or PGG had significantly reduced tumor burden (\*\**P* < 0.01 and \*\*\**P* < 0.001, respectively). The PGG+Ola combination treatment achieved the most pronounced anti-tumor effect and almost completely arrested tumor growth, resulting in significantly smaller tumors versus the Ola monotherapy group (*P* < 0.05). Gross evaluation of the excised tumors at the experimental endpoint (day 16) confirmed that the combination regimen yielded the most profound reduction in physical tumor size across all cohorts (**Figure 6B**). In alignment with these morphological observations, the average tumor weight in the PGG+Ola group was drastically minimized to 0.05 g, presenting a sharp contrast to the 0.22 g and 0.18 g recorded in the Ola and PGG monotherapy arms, respectively (**Figure 6C**). According to these in vivo findings, the combination of PGG and Ola exerts a superior anti-tumor efficacy, robustly arresting tumor progression and alleviating overall tumor burden.

**Figure 6.**
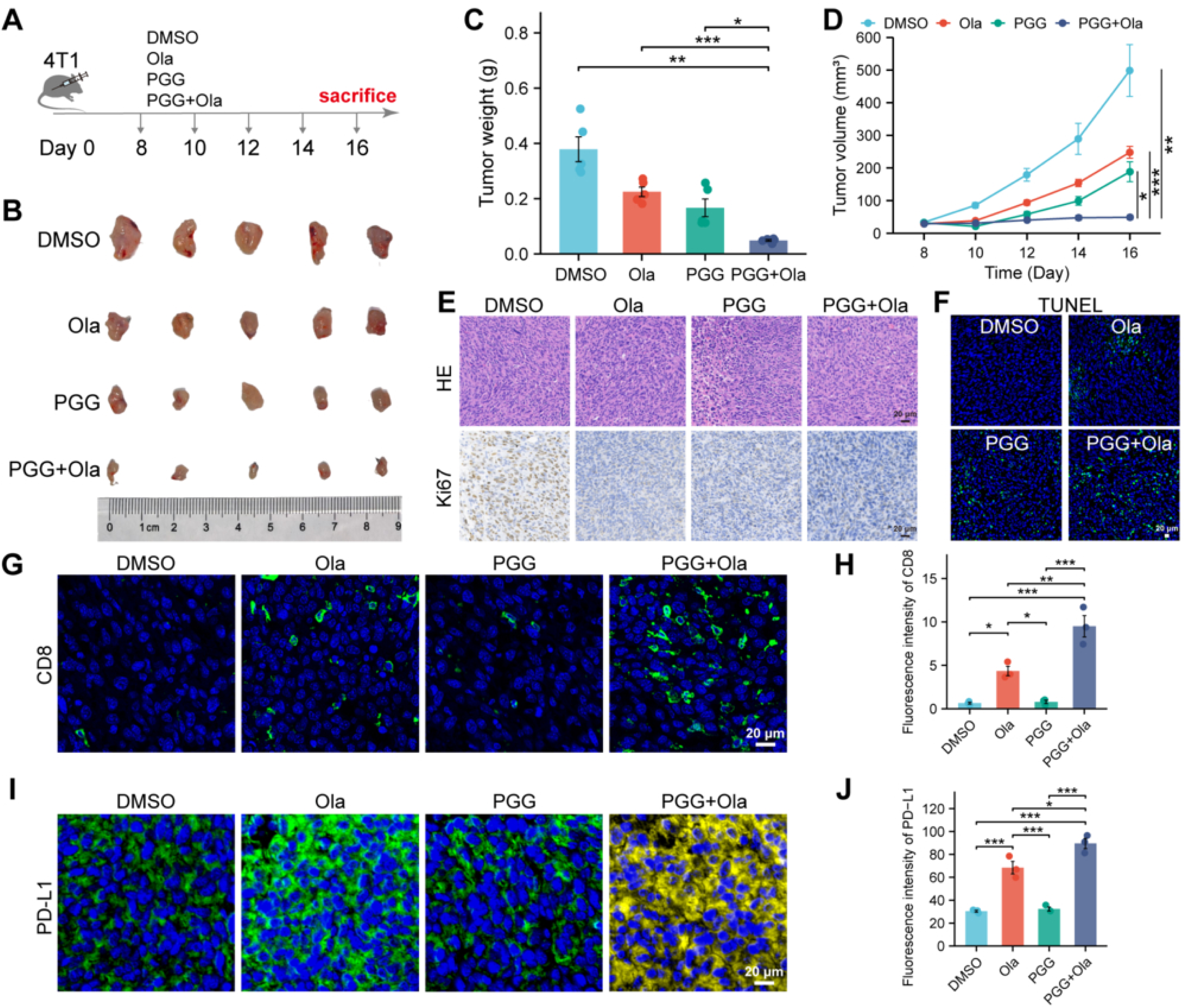
Immunomodulatory and anti-tumor effects of the PGG and Ola combination in vivo. **(A)** Schematic illustration of the establishment of the 4T1 subcutaneous syngeneic tumor model and treatment regimens. **(B)** Representative photographs of excised tumors harvested from the mice in different treatment groups on day 16 (n=5). **(C)** Mean tumor weights in the indicated groups at the experimental endpoint. **(D)** Dynamic tumor growth curves demonstrating changes in tumor volume in the indicated groups over the 16-day experimental cycle. **(E)** Representative H&E staining (top panel) and Ki-67 immunostaining (bottom panel) images of tumor sections from the indicated groups (scale bars = 20 *μ*m). **(F)** Representative fluorescence images of TUNEL staining showing apoptosis in the tumor tissues from indicated groups (green: TUNEL-positive apoptotic cells; blue: DAPI-stained nuclei; scale bar = 20 *μ*m). **(G)** Representative immunofluorescence images showing infiltration of CD8^+^ T cells (green) within the TME in the indicated groups (blue: DAPI; scale bar = 20 *μ*m).**(H)** Quantification of the MFIs of CD8 expression in the tumor tissues based on ImageJ analysis. **(I)** Representative immunofluorescence images illustrating the expression levels and distribution of PD-L1 (green/yellow) in the indicated groups (blue: DAPI; scale bar = 20 *μ*m). **(J)** Quantification of the MFIs of PD-L1 expression in the indicated groups. Data are in the format of mean±SEM. \**P* < 0.05, \*\**P* < 0.01, \*\*\**P* < 0.001.

With the objective of elucidating the anti-tumor effect of PGG+Ola at the cellular level, we analyzed the tumor tissues from each group. H&E staining revealed varying degrees of tissue necrosis across all treatment groups versus the DMSO control group. The proliferative capacity of the tumor cells was assessed by Ki-67 immunostaining. The DMSO control group possessed the highest percentage of Ki-67+ cells, indicating robust proliferation of tumor cells. While there were less Ki-67-positive cells after separate PGG and Ola treatment, the PGG+Ola group displayed the lowest Ki-67 expression level, indicating that this combination regimen effectively inhibited the proliferation of 4T1 cells in vivo (**Figure 6E**). Furthermore, TUNEL staining was employed to evaluate apoptosis in the tumor tissues (**Figure 6F**). In contrast to the sparse and sporadic apoptotic signals observed in the control group, both PGG and Ola induced noticeable apoptosis in the tumor cells. The strongest TUNEL signals were detected in the PGG+Ola combination group, indicating that PGG and Ola achieved significant tumor suppression by triggering massive apoptosis in a cooperative manner.

To investigate whether PGG-mediated inhibition of DNA damage repair could trigger an anti-tumor immune response, we evaluated immune infiltration and PD-L1 expression in the tumor tissues across all groups. Both the PGG and Ola monotherapy groups exhibited moderately higher intra-tumor infiltration of CD8^+^ T cells versus the DMSO control group. The PGG+Ola combination group demonstrated the densest infiltration of CD8^+^ T cells, as indicated by the remarkably stronger fluorescence intensity versus the control and monotherapy groups (**Figure 6G**, **6H**).

This finding suggests that the blockade of PALB2-BRCA2 binding by PGG can effectively enhance tumor immunogenicity, thereby promoting cytotoxic T cells to be recruited into the TME. Besides, the combination treatment significantly increased PD-L1 expression on the tumor cells versus the control and monotherapy groups (**Figure 6I**, **6J**). This in vivo phenotype conformed to our in vitro results, and confirmed that the combination therapy excels in activating the STING pathway and upregulating PD-L1.

To exploit the compensatory upregulation of PD-L1 induced by PGG+Ola dual therapy, we evaluated a triple combination regimen incorporating an anti-PD-L1 (aPD-L1) antibody in the syngeneic 4T1 subcutaneous tumor model (**Figure 7A**). While aPD-L1 monotherapy or PGG+Ola dual therapy achieved moderate tumor inhibition, the triple combination regimen achieved near-complete tumor growth arrest, demonstrating significantly abated growth kinetics (**Figure 7C**) and minimized endpoint tumor burden compared with all other cohorts (**Figure 7B, D**). Consistent with the alleviated tumor burden, H&E and Ki-67 staining revealed extensive intratumoral necrosis and profoundly suppressed cellular proliferation within the triple-therapy cohort (**Figure 7E**). Consistently, TUNEL assays demonstrated a robust surge in apoptotic signals, confirming widespread cell death in vivo (**Figure 7F**). Immunofluorescence profiling of the tumor microenvironment (TME) revealed that the triple combination therapy triggered massive intratumoral infiltration of CD8^+^ T cells, with the MFI significantly exceeding that of the PGG+Ola dual-therapy arm (**Figure 7G, H**). Concurrently, aPD-L1 administration effectively targeted and neutralized the surface-accessible PD-L1 on tumor cells (**Figure 7G, I**). Collectively, these findings demonstrate that PD-L1 blockade successfully bypasses the compensatory checkpoint barrier erected by PGG+Ola dual therapy, maximizing tumor eradication through the concerted action of immune reactivation and DNA damage accumulation.

**Figure 7.**
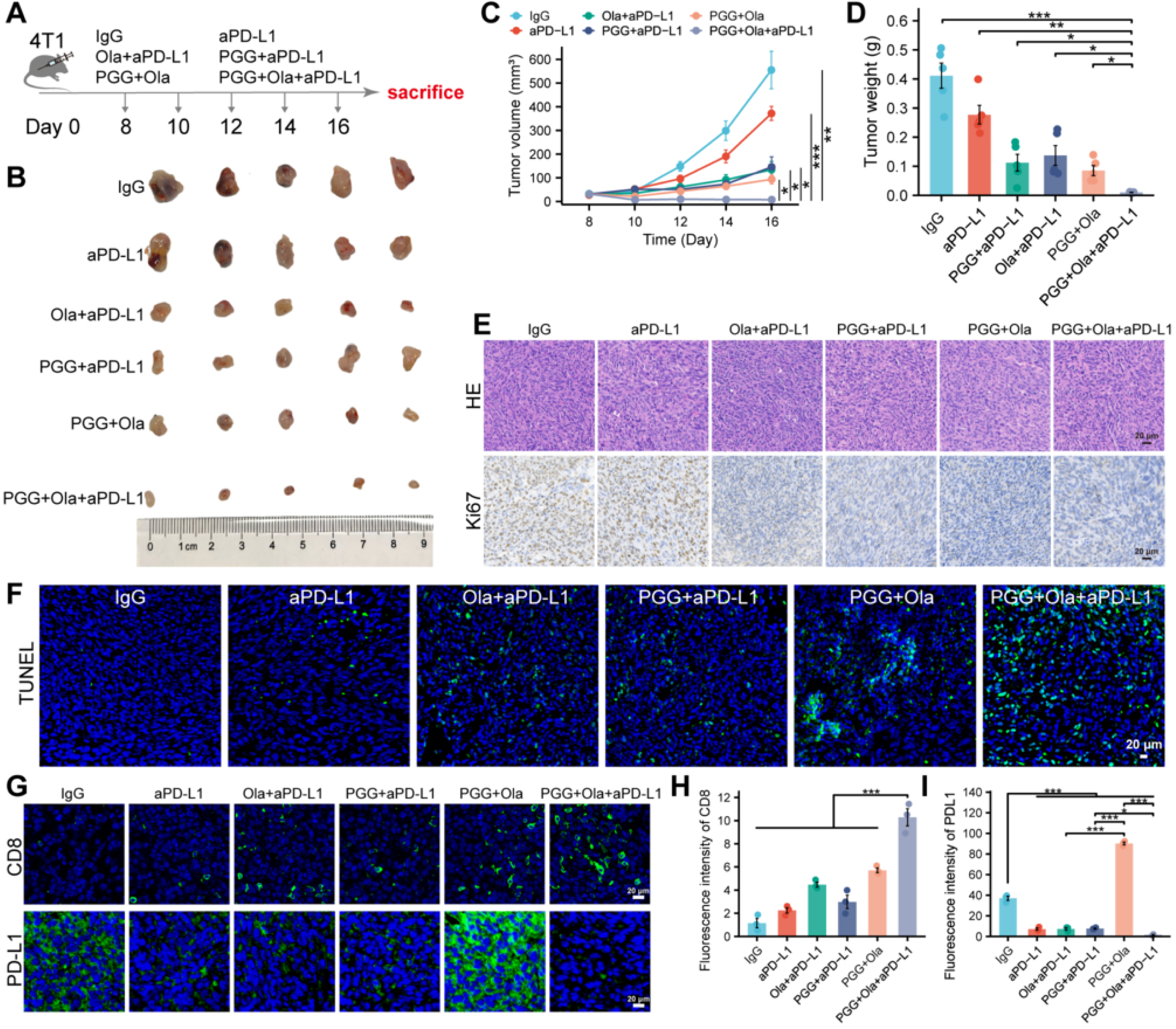
PGG and Ola cooperatively suppressed tumor growth and modulated immune responses with anti-PD-L1 therapy. **(A)** Schematic illustration of the establishment of the 4T1 subcutaneous syngeneic tumor model and the treatment regimens. **(B)** Representative photographs of excised tumors harvested from the mice in varying treatment groups on day 16 (n=5). **(C)** Dynamic tumor growth curves demonstrating the varying tumor volume during the 16-day evaluation period. **(D)** Mean tumor weights in the indicated groups on day 16. **(E)** Representative H&E staining (top panel) and Ki-67 immunostaining (bottom panel) images of tumor tissues from the indicated groups (scale bars = 20 *μ*m). **(F)** Representative fluorescence images of TUNEL staining showing apoptosis in the tumor tissues from indicated groups (green: TUNEL-positive apoptotic cells; blue: DAPI-stained nuclei; scale bar = 20 *μ*m). **(G)** Representative immunofluorescence images showing CD8^+^ T cells (green) infiltrating within the TME in the indicated groups (blue: DAPI; scale bar = 20 *μ*m). Representative immunofluorescence images illustrating the expression levels and distribution of PD-L1 (green) in the indicated groups (blue: DAPI; scale bar = 20 *μ*m). **(H)** Quantification of the MFIs of CD8 expression in the tumor tissues based on ImageJ analysis. **(I)** Quantification of the MFIs of PD-L1 expression in the indicated groups. Data are in the format of mean±SEM. \**P* < 0.05, \*\**P* < 0.01, \*\*\**P* < 0.001.

## Discussion

TNBC remains the most challenging BC subtype relying on its high aggressiveness, lack of defined therapeutic targets, and poor prognosis. Although PARPi like Ola have transformed the treatment paradigm for TNBC through synthetic lethality, their clinical utility is severely limited by drug resistance and low mutation frequency in patients. According to our finding, PGG, a specific small-molecule inhibitor of the PALB2-BRCA2 interaction, can effectively induce an HRD state in TNBC cells. PGG markedly sensitized the BRCA-wild-type TNBC cells to Ola, which not only suppressed short-term proliferation, but also attenuated their long-term clonogenic potential, and inhibited tumor growth in vivo. Furthermore, the PGG+Ola-mediated restriction of TNBC cell migration and invasion in vitro carries profound clinical significance, directly addressing the early systemic dissemination and high motility that characterize this aggressive malignancy. Consequently, these findings underscore that pharmacological disruption of the PALB2-BRCA2 axis serves as a powerful strategy to counteract the metastatic propensity of BRCA-wild-type TNBC cells.

Programmed death cascades are accompanied by damage-associated molecular patterns being well released, which can convert an immunosuppressive TME into an immunologically active milieu. Furthermore, recent studies have shown that ICD induced by PANoptosis can effectively reverse tumor immune evasion [30]. Consistent with this, the combination of PGG and Ola aggravated intracellular ROS accumulation and mitochondrial damage, which directly precipitated mitochondrial apoptosis and drove the subsequent release of dangerous signals, thereby exhibiting characteristic features of an ICD-like phenotype.

The accumulation of DNA lesions effectively triggers the cytosolic innate immune surveillance network [31]. The process of dsDNA leaking into the cytoplasm is accompanied by the activation of the cGAS-STING pathway, facilitating STING and IRF3 to be phosphorylated, subsequently driving cytotoxic CD8^+^ T cell recruitment into the TME [32]. In our study, genetic loss-of-function assays via siRNA-mediated knockdown of STING (siSTING) confirmed that the STING pathway activation mediated the combined therapeutic effect of PGG and Ola. However, STING activation also upregulates PD-L1 on TNBC cell surface, aiding immune evasion [33]. Notably, PD-L1 expression remained elevated even under the pharmacological blockade of STING, which suggests that non-canonical bypass pathways triggered by DNA damage may sustain the immune checkpoint, thereby suppressing T cell-mediated cytotoxicity. Therefore, we introduced aPD-L1 into the treatment regimen to block this adaptive feedback loop; the triple combination therapy (PGG+Ola+aPD-L1) achieved near-complete tumor eradication by triggering a surge of apoptotic cascades, and inducing massive intra-tumoral infiltration of CD8^+^ T cells.

Beyond its established role in disrupting intracellular DNA repair, our bioinformatic modeling offers an intriguing theoretical rationale for the potential involvement of PGG in stromal remodeling. Enrichment analysis revealed that the overlapping targets of PGG and TNBC cluster prominently within pathways governing “extracellular matrix disassembly,” “degradation of the extracellular matrix,” and “complement and coagulation cascades”. The dense, desmoplastic stroma of TNBC is characterized by excessive collagen deposition and cross-linking, which increase matrix stiffness and compress the intra-tumoral blood vessels. Vascular compression leads to high interstitial fluid pressure (IFP) and severe hypoxia, which physically impede the influx of therapeutic agents and TILs [34]. Based on our bioinformatic modeling, PGG is predicted to target pathways involved in ECM degradation in the tumor stroma, hypothetically influencing matrix stiffness, intra-tumoral IFP, and vascular normalization [35]. The normalized vessels restore efficient perfusion, which enhances the cGAS-STING-driven chemokine gradients to stimulate CD8^+^ T cells to be recruited into the stroma [36]. Furthermore, alleviating mechanical stiffness directly removes the physical triggers that stimulate epithelial-mesenchymal transition, thereby inhibiting tumor invasion and metastasis [37].

From a mechanobiological perspective, PGG-induced structural relaxation of the ECM can also regulate the stromal cells, particularly the cancer-associated fibroblasts (CAFs). According to recent single-cell RNA sequencing and spatial transcriptomics research, the mechanical properties of ECM dictate fibroblast heterogeneity and activation states [38]. A rigid, high-stiffness microenvironment continuously provides mechanical cues that sustain the activation of pro-tumorigenic CAFs that synthesize collagen and secrete immunosuppressive cytokines, thereby reinforcing the cold tumor state [39]. Alleviating physical tension in the stroma alters the gene expression profiles of these fibroblasts, reverting them from an aggressive, matrix-depositing phenotype to an immunologically quiescent state [40]. This mechanobiological resetting not only slows down the self-perpetuating cycle of desmoplasia but also reverses the immunosuppressive niche maintained by CAFs [41]. Our findings suggest that PGG can effectively disrupt the mechanotransduction feedback loop in tumor stroma by softening the ECM. Thus, PGG acts as a multi-functional therapeutic agent that coordinates intracellular synthetic lethality, innate immune sensing activation, and stromal normalization to achieve maximal therapeutic efficacy against aggressive breast malignancies. Although further experimental validations, such as collagen staining and immunofluorescence of vascular markers, are warranted to visually confirm these structural alterations, our current findings offer a distinct mechanobiological perspective on how PGG multi-functionally reshapes the desmoplastic stroma.

## Conclusion

PGG induces an HRD state in TNBC cells by disrupting the PALB2-BRCA2 interaction, thereby exerting a potent synthetic lethal effect when combined with Ola. Mechanistically, the combination of PGG and the PARPi aggravated mitochondrial damage and ROS accumulation, and increased the accumulation of unrepaired DNA lesions, which culminated in robust apoptosis and the emission of ICD-associated DAMPs. Concurrently, the cytosolic leakage of dsDNA activated the STING-IRF3 axis and promoted cytotoxic CD8^+^ T cells to be infiltrated into the TME. Since this pathway concurrently induces adaptive PD-L1 upregulation, we also incorporated aPD-L1 into the treatment regimen to neutralize the checkpoint barrier. The triple combination therapy achieved near-complete tumor eradication in vivo. Furthermore, bioinformatics analyses propose a tentative role for PGG in the hypothetical regulation of the desmoplastic stroma. The predicted targets suggest that PGG might participate in the degradation of the ECM, which remains to be experimentally validated regarding its physical impacts on matrix stiffness, vascular normalization, and CAF phenotypes. Overall, PGG serves as a multi-functional therapeutic agent integrating intracellular synthetic lethality and cytosolic innate immune sensing activation, with a promising bioinformatic hypothesis for prospective microenvironmental remodeling. Collectively, this study establishes PGG as a multi-functional agent integrating intracellular synthetic lethality and cytosolic innate immune activation to overcome PARPi resistance, while offering a preliminary bioinformatic rationale for prospective microenvironmental remodeling in triple-negative breast cancer.

## Data Availability Statement

All data generated or analyzed in this study can be obtained in this published article. The public datasets analyzed during this study are openly available in [the HERB database] at http://herb.ac.cn and [GeneCards database] at https://www.genecards.org.

## Conflict of Interest Statement

The authors declare no conflicts of interest.

## Author Contributions

S.L: original draft, data curation; X.L: original draft, data curation; X.Y: original draft, data curation; M.X: original draft, formal analysis; T.F: original draft, formal analysis; J.Y:writing-review and editing, funding; J.Z: writing-review and editing, funding, supervision. All authors were involved in drafting and revising the manuscript.

## Acknowledgements

Gemini (accessed March 2026) was used to reorganize the Discussion section for clarity and flow. All AI-assisted text was reviewed and revised by the authors to ensure accuracy and clarity of meaning.

## Fundings

The Natural Science Foundation of Hunan Province [grant number 2024JJ9320], the Chronic Disease Management Research Project of the National Health Commission Capacity Building and Continuing Education Center [grant number GWJJMB202510022060], and Hunan Provincial Health High-Level Talent Scientific Research Project (grant number: R2023140)

## Ethics Approval

The Experimental Animal Ethics Committee of Hunan Normal University granted approval for all in vivo experiments (Approval No. [2024] 97). All animal handling, care, and euthanasia procedures strictly followed the institutional guidelines for laboratory animal welfare and protection to minimize animal suffering.

## Notes

### Competing Interest Statement

The authors have declared no competing interest.

