## Supplemental Material for "PGG Potentiates Olaparib-Induced DNA Damage and STING-Associated Immunogenic Signaling to Enhance PD-L1 Blockade in Triple-Negative Breast Cancer"

**Supplementary Information**

**Supplementary Note: Rationale for the Selection of in vitro PGG and Olaparib Concentrations**

The working concentrations of 1,2,3,4,6-penta-O-galloyl-β-D-glucose (PGG, 1 μM) and olaparib (Ola, 10 μM) for in vitro cellular assays (including CCK-8, colony formation, ROS/MMP detection, and Western blotting) were determined based on our group’s previously published dose-response characterization and mechanistic validations. (J. Zeng, J. Han, Z. Liu, M. Yu, H. Li, and J. Yu, "**Pentagalloylglucose Disrupts the PALB2-BRCA2 Interaction and Potentiates Tumor Sensitivity to PARP Inhibitor and Radiotherapy**." *Cancer Letters* 546 (2022): 215851, <https://doi.org/10.1016/j.canlet.2022.215851>.)

**1. PGG (1 μM):** In our previous study, co-immunoprecipitation assays demonstrated that 1 μM PGG effectively disrupts the endogenous PALB2–BRCA2 protein-protein interaction (**Fig. 1a**). Moreover, the DR-GFP reporter assay validated that 1 μM PGG significantly suppresses homologous recombination (HR) repair efficiency by over 50% (**Fig. 2b**). At this concentration, PGG exerts minimal cytotoxicity as a monotherapy (MDA-MB-231 monotherapy IC_50_ = 3.24 μM, **Fig. 3a**) and shows negligible toxicity in normal human mammary epithelial cells MCF-10A (**Fig. 4a**), serving as an ideal pharmacological dose to induce homologous recombination deficiency (HRD) without non-specific cytotoxicity.

**2. Olaparib (10 μM):** As established previously, the baseline IC_50_ of olaparib monotherapy in BRCA-proficient MDA-MB-231 cells was approximately 9.7 μM (9704 nM) (**Fig. 4b**). Thus, 10 μM represents a physiologically and experimentally relevant near-IC_50_ baseline concentration for testing combination-induced synthetic lethality in BRCA-wild-type TNBC cells.

**3. Synergistic Potentiation:** Our prior findings demonstrated that 1 μM PGG synergizes robustly with olaparib in MDA-MB-231 cells, reducing the olaparib IC_50_ by more than 10-fold (from 9704 nM down to 907.7 nM; **Fig. 4b**) and suppressing colony formation (**Fig. 4e**). Therefore, the combination of 1 μM PGG and 10 μM olaparib provides a well-defined therapeutic window to capture synthetic lethality, oxidative stress, mitochondrial collapse, and subsequent cGAS-STING-mediated immune activation.

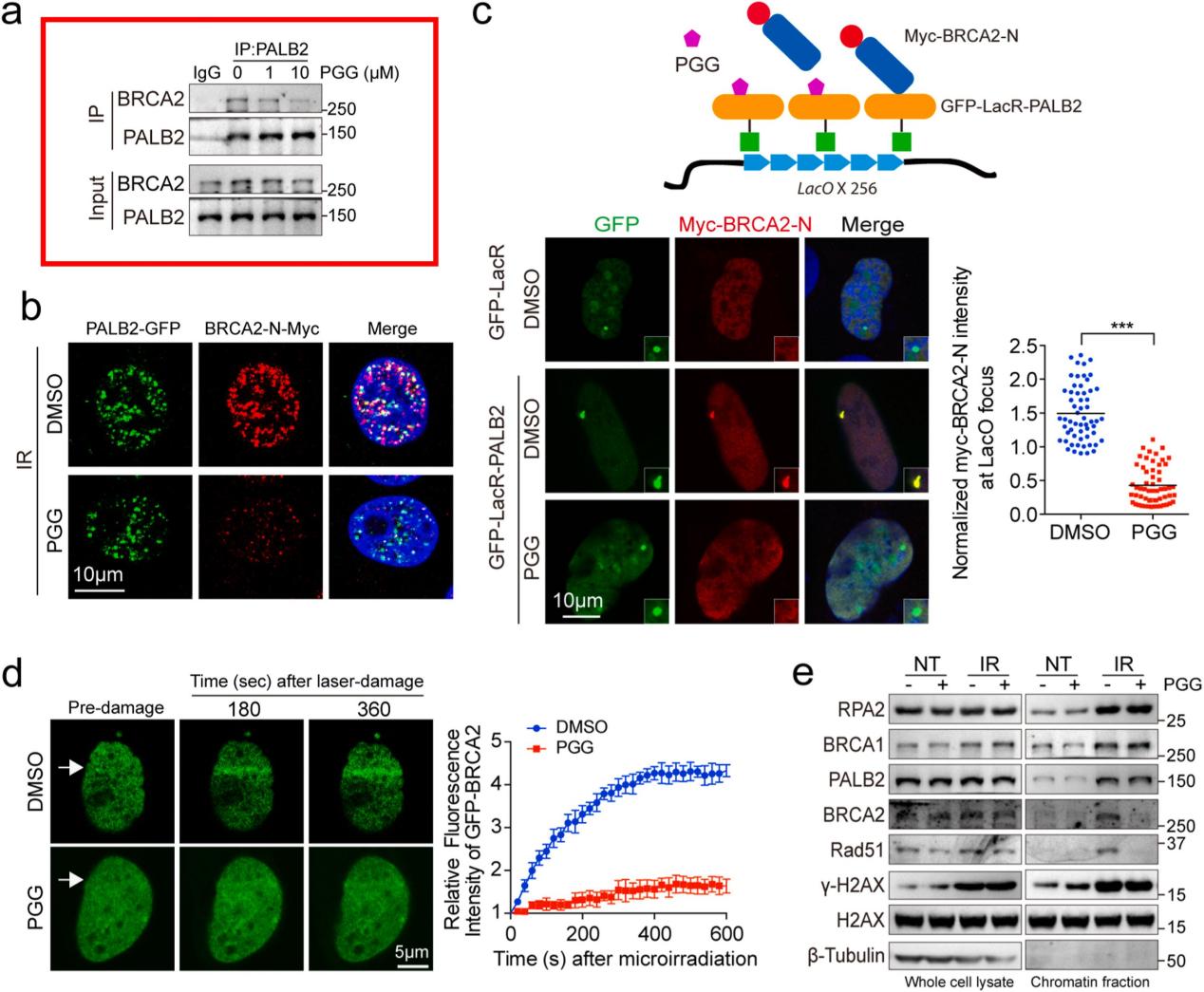

**Figure 1. PGG disrupts the PALB2-BRCA2 interaction.** (a) Co-immunoprecipitation assay of PALB2 and BRCA2 from lysates of U2OS cells treated with DMSO or 1 μM or 10 μM PGG 30 min before ionizing radiation (IR, 10 Gy). Western blotting was performed with anti-PALB2 and anti-BRCA2 4 h after IR.

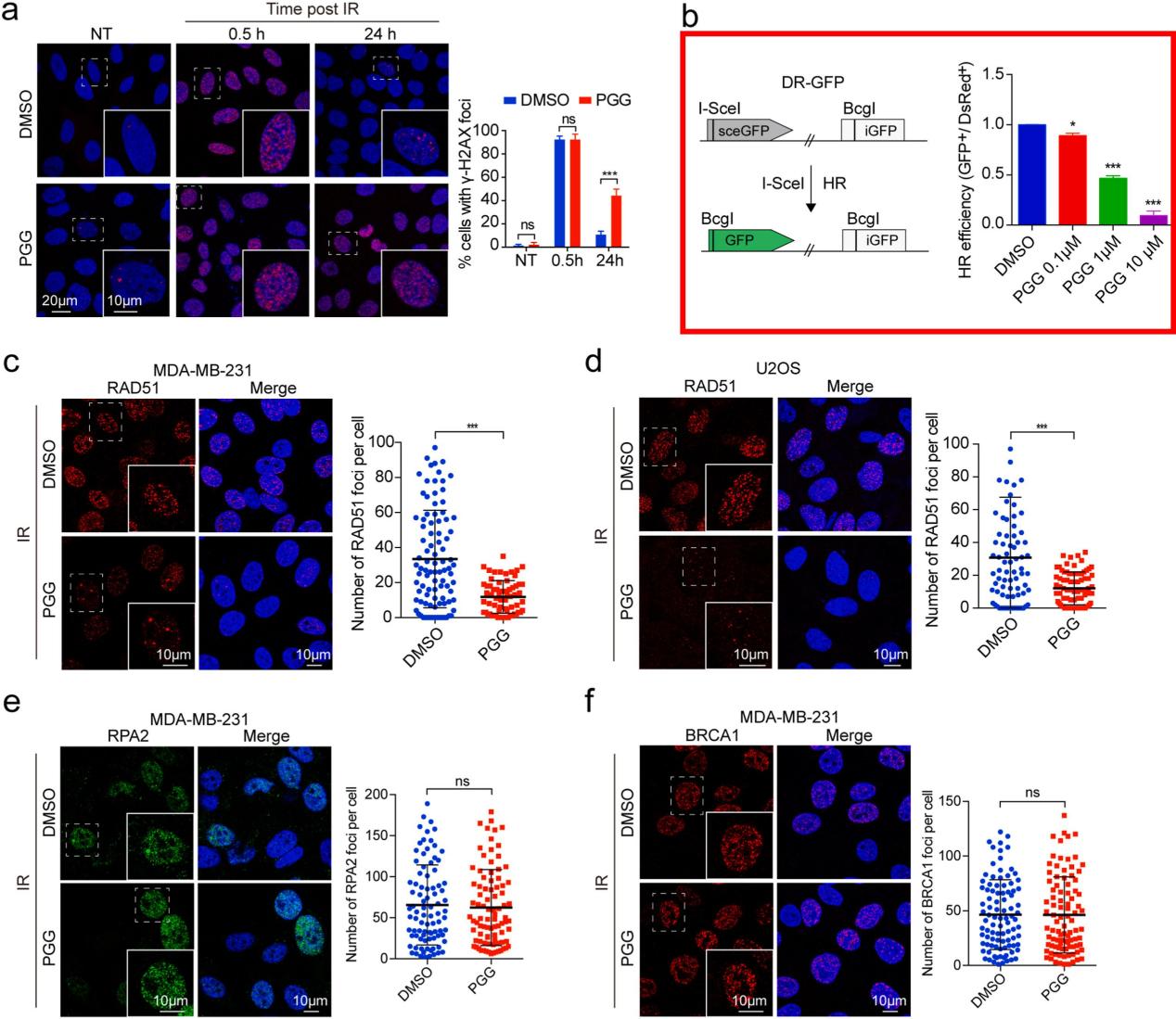

**Figure 2. PGG inhibits HR-mediated DSB repair.** (b) Scheme of the DR-GFP reporter system to assess HR efficacy. U2OS DR-GFP cells were treated with DMSO and increasing concentrations of PGG (0.1, 1, and 10 μM) for 12 h after transfection with I-SceI plasmid. GFP-positive cells were detected by flow cytometry. Data are presented as the mean ± SD (n = 3 independent experiments; paired t-test; **p* < 0.05; ****p* < 0.001).

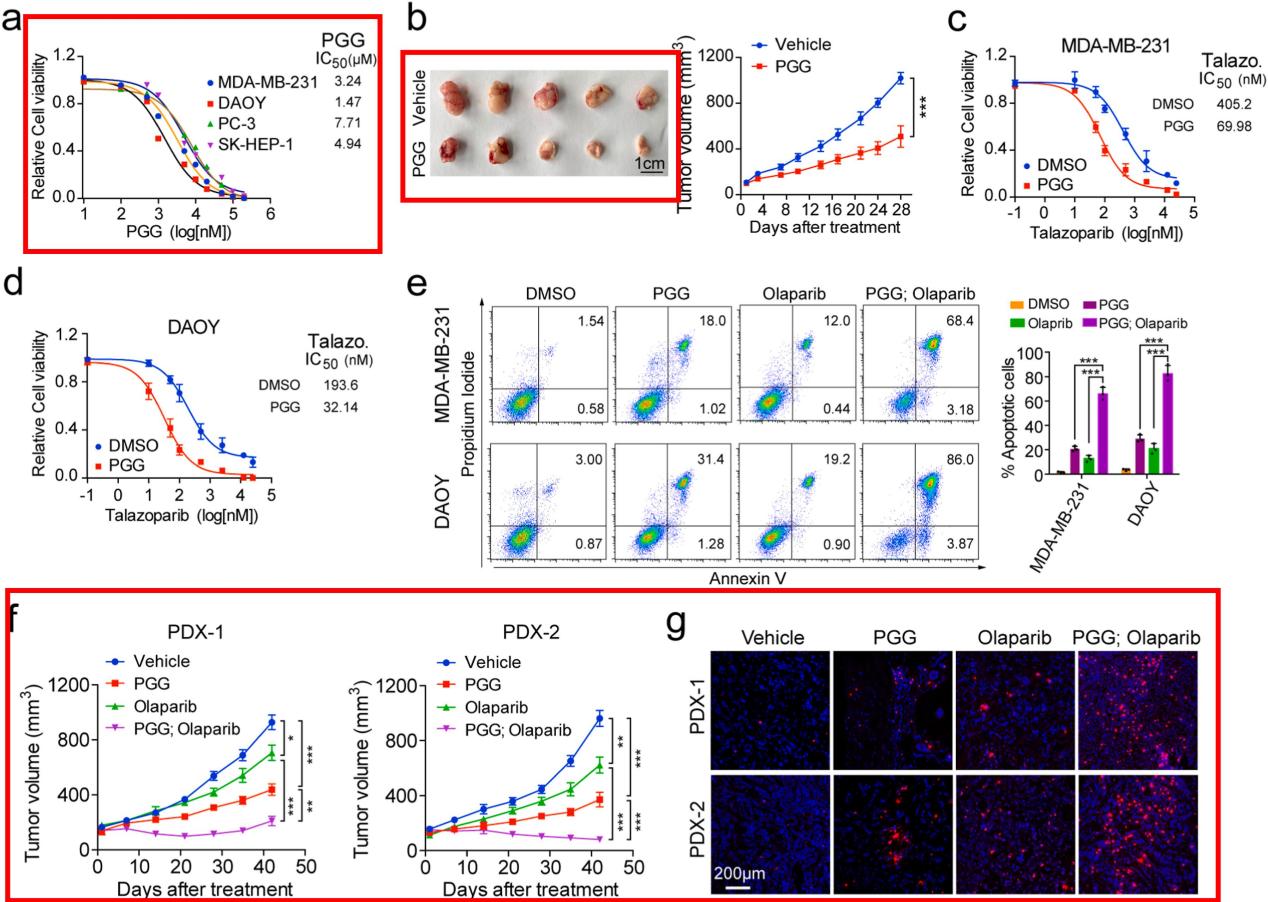

**Figure 3. PGG sensitizes tumor cells to PARP inhibitor.** (a) Viability curves of the indicated cell lines in response to PGG. Cells were treated for 5 days, and cell viability was measured using CellTiter-Glo. Data represent the mean ± SD (n = 3 independent experiments). The IC_50_ was calculated using GraphPad Prism 6 software.

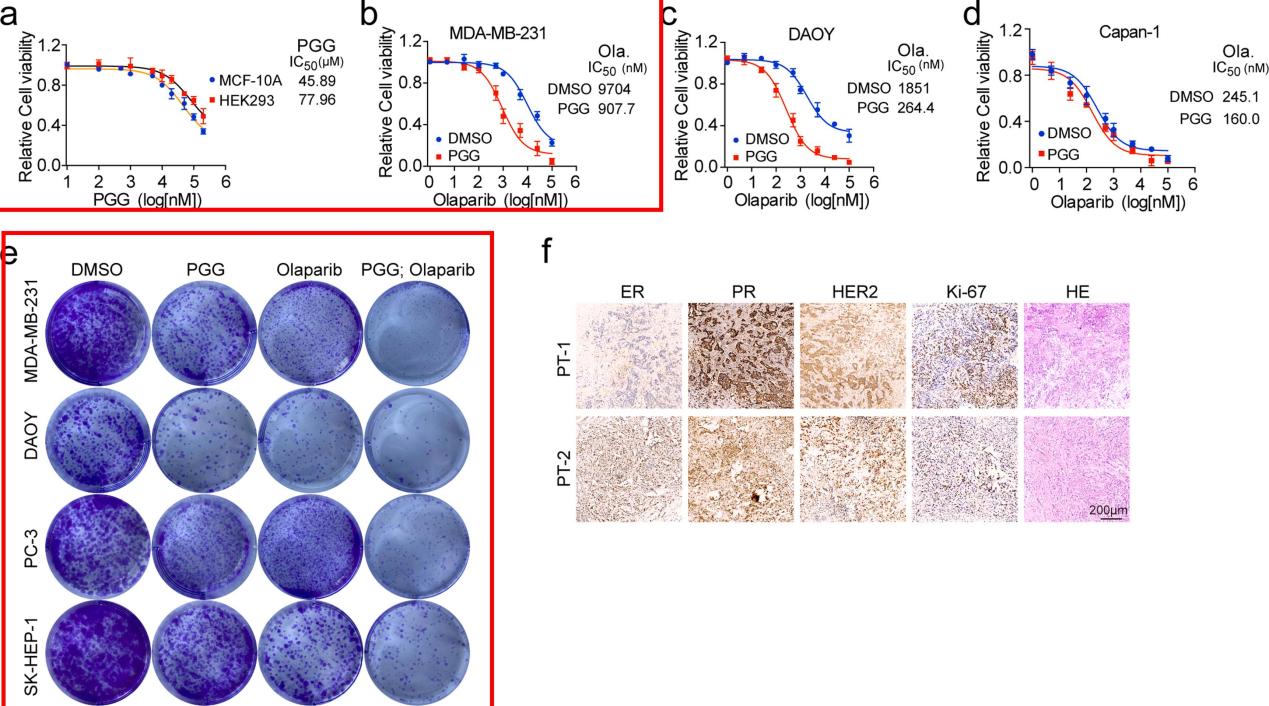

**Figure 4.**  (a) Viability curves of MCF-10A and HEK293 cells in response to PGG. Cells were treated for 5 days, and the cell viability was measured using CellTiter-Glo. Data represent the mean ± SD (n = 3 independent experiments). The IC50 was calculated using GraphPad Prism 6 software. (b-d) Olaparib dose-response curves of MDA-MB-231 (b), DAOY (c), and Capan-1 (d) cells treated with the indicated concentrations of olaparib and PGG (1 μM) or DMSO. The cells were treated for 5 days, and cell viability was measured using CellTiter-Glo. Data represent the mean ± SD (n = 3 independent experiments). The IC_50_ was calculated using GraphPad Prism 6 software. (e) Effect of combinational treatment with PGG and olaparib on cell clonogenicity. The indicated cell lines were treated with PGG (1 μM), olaparib (1 μM), or the combination for 2 weeks. The colonies were fixed and stained with crystal violet.

**Supplementary Note: Rationale for in vivo Dosing Regimens of PGG and Olaparib**

The in vivo dosages and administration routes of 1,2,3,4,6-penta-O-galloyl-β-D-glucose (PGG, 10 mg/kg, i.v., q.2.d.) and olaparib (Ola, 50 mg/kg, i.p., q.d.) in the syngeneic 4T1 tumor model were selected based on the pharmacokinetic profiles, safety tolerability, and antitumor efficacy previously established by our research group. (J. Zeng, J. Han, Z. Liu, M. Yu, H. Li, and J. Yu, "**Pentagalloylglucose Disrupts the PALB2-BRCA2 Interaction and Potentiates Tumor Sensitivity to PARP Inhibitor and Radiotherapy**." *Cancer Letters* 546 (2022): 215851, <https://doi.org/10.1016/j.canlet.2022.215851>.)

**1. PGG (10 mg/kg, i.v.) Administration and Pharmacokinetics:** Our prior pharmacokinetic (PK) investigation in mice demonstrated that intravenous administration of PGG at 10 mg/kg ensures rapid systemic distribution and favorable plasma bioavailability (**Table 1**). Conversely, oral administration suffered from extensive intestinal degradation due to ester bond hydrolysis. Furthermore, intravenous delivery of 10 mg/kg PGG exhibited demonstrable in vivo tumor growth delay in MDA-MB-231 xenografts (**Fig. 3b**) without observable adverse systemic toxicity or significant body weight loss.

**2. Olaparib (50 mg/kg) Baseline Dosage:** Olaparib at 50 mg/kg represents a standard, biologically active in vivo dose widely utilized in preclinical models. In our previous study, 50 mg/kg olaparib monotherapy produced moderate growth-retarding effects in BRCA-wild-type patient-derived xenografts (PDX-1 and PDX-2) (**Fig. 3f**), thereby providing an optimal baseline window to evaluate synergistic synthetic lethality in vivo.

**3. In Vivo Synergistic Efficacy and DNA Damage:** As established in our 2022 study, combining 10 mg/kg PGG with 50 mg/kg olaparib resulted in superior therapeutic synergy, leading to profound tumor arrest in PDX-1 and marked tumor regression in PDX-2 (**Fig. 3f** ). Histological analysis further revealed that this dual regimen provoked maximal accumulation of unrepaired DNA double-strand breaks (γ-H2AX foci) in excised tumor tissues (**Fig. 3g**). Hence, this optimized regimen was directly translated into the current immunocompetent syngeneic model to investigate the downstream cGAS-STING innate immune response and combinatorial PD-L1 blockade.

| **Table 1. Pharmacokinetics of PGG in C57BL/6 mice intravenously and orally** | | | | | |
| --- | --- | --- | --- | --- | --- |
|  | Table (1). Plasma concentration-time profiles of PGG in C57BL/6 mice after intravenous administration (10mg/kg) (n=3) | | | | |
|  |  | Time | PGG Concentration in plasma（ng/mL） | | |
|  | （h） | 1 | 2 | 3 | Average |
|  | 0.08 | 13103.3 | 5404.77 | 6378.21 | 5891.49 |
|  | 0.25 | 1892.25 | 1333.06 | 1604.31 | 1609.87 |
|  | 1 | 788.12 | 769.23 | 823.29 | 793.55 |
|  | 2 | 571.11 | 345.4 | 509.45 | 475.32 |
|  | 4 | 263.59 | 289.5 | 416.23 | 323.11 |
|  | 8 | 189.02 | 211.79 | 206.94 | 202.58 |
|  | 24 | 233.18 | 189.84 | 194.77 | 205.93 |
|  | Note: The data marked with red has poor paralelism, so it is excluded when calculating pharmacokinetic parameters. | | | | |
|  | Table (2). Pharmacokinetic parameters of PGG in C57BL/6 mice after intravenous administration (10mg/kg) (n=3) | | | |  |
|  |  | Parameter | Units | i.v. 10 mg/kg | |
|  | AUC_(0-t)_ | h·µg/L | 7738.37±594.56 | |  |
|  | AUC_(0-∞)_ | h·µg/L | 12997.43±2292.57 | |  |
|  | t_1/2_ | h | 18.22±10.02 | |  |
|  | V | L/kg | 19.40±7.07 | |  |
|  | CL | L/h/kg | 0.78±0.13 | |  |
|  | MRT_(0-∞)_ | h | 8.04±0.83 | |  |
|  | Table (3). Brain concentration-time profiles of PGG in C57BL/6 mice after intravenous administration (10mg/kg) (n=3) | | | | |
|  |  | Time | PGG Concentration in brain（ng/g brain） | | |
|  | （h） | 1 | 2 | 3 | Average |
|  | 0.08 | ND | ND | 76.76 | --- |
|  | 0.25 | ND | ND | ND | --- |
|  | 1 | ND | ND | ND | --- |
|  | 2 | ND | ND | ND | --- |
|  | 4 | 101.44 | 66.08 | ND | 83.76 |
|  | 8 | 93.28 | 109.16 | 83.94 | 95.46 |
|  | 24 | ND | 82.41 | ND | --- |
|  | Note: ND means lower than the limit of quantification, not detected. | | | | |
